# The mouse lung early cellular innate immune response is not sufficient to control fungal infection with Cryptococcus neoformans

**DOI:** 10.1101/679274

**Authors:** Jacob Rudman, Helen Maria Marriott, Leo M. Carlin, Simon Andrew Johnston

## Abstract

*Cryptococcus neoformans* causes life-threatening infection in the immunocompromised. This and other opportunistic pathogens are an increasing threat as immunosuppression increases globally. To counter antibiotic resistance, there is precedent for developing immune enhancing therapy. However, our understanding of how immunocompetent patients resolve these infections is poor as opportunistic infections typically resolve subclinically. Because this has led to a lack of clinical data, we rely on animal models. Current *in vivo* infection models either lack mammalian immunity or are not compatible with long term high content imaging required to model the complexities of human host-pathogen interactions. Therefore, we have developed an *ex vivo* murine precision cut lung slice (PCLS) model to understand innate immunity in cryptococcosis. C57BL/6 mice were sacrificed 0 or 24 hours post infection with *KN99α* cryptococci. Lungs were inflated with 37°C agarose, 300μm thick PCLS were prepared on a vibratome and imaged by confocal or wide-field fluorescence microscopy. Using PCLS and immunofluorescence, we demonstrate cryptococcal replication and clearance rates are balanced over the first 24 hours of infection. Cell-mediated immunity is alveolar macrophage centric, although alveolar macrophages demonstrate limited phagocytosis of cryptococci and enable intracellular cryptococcal replication. *Cryptococcus neoformans* responded to the lung environment by forming enlarged cells, although these were not large enough to be titan cells. To further understand cryptococcal proliferation *in vivo*, we also infected animals with *plb1* mutant *Cryptococcus neoformans* that has been shown to exhibit proliferation defects *in vivo*. We found no difference in fungal burden with *plb1* infected animals 24 hours post infection, but observed significantly larger fungal cells and no incidences of phagocytosis. Thus, the PCLS model can be used to assess the lung immune response early in cryptococcal infection, demonstrating that resident lung macrophages cannot control cryptococcal infection and offer an intracellular niche for *Cryptococcus neoformans* growth.

## Introduction

Opportunistic infections are a class of infections that are readily cleared by healthy individuals, yet in immunosuppressed patients can be life threatening. In these patients, balancing immunosuppression and infection clearance complicates treatment. When combined with an increasing incidence of antimicrobial resistance (1), there is a clear need to better understand normal immunity to opportunistic infections in an effort to develop new therapies, particularly those that may be immunomodulatory (2, 3).

*Cryptococcus neoformans* is an example of an often-fatal opportunistic pathogen where the physiological immune response is still poorly understood. Cryptococcal infection is one of the leading causes of mortality in HIV/AIDS patients (15%), estimated to cause over 200,000 infections globally a year and more than 180,000 deaths (4). *C. neoformans* has evolved multiple virulence mechanisms related to an ecological niche in the soil and tree saprophyte in response to natural predators such as amoebae that have allowed this fungus to cause disease (5, 6). These include a polysaccharide capsule, production of oxygen scavenging melanin and the ability to modulate host immunity (7–10). The majority of cryptococcosis patients present late in disease, following dissemination of infection to the CNS – meaning most experimental and clinical research is focussed on these later stages (14, 15, 16, 17). Once diagnosed, the current gold standard of treatment is a combination of antifungal medications, all of which have major issues including limited availability, widespread resistance and hepatotoxicity (15, 16). Therefore, cryptococcosis is a perfect example of an opportunistic infection where better, alternative treatments are needed, informed by a greater understanding of the immune response. Therefore, we sought to develop an experimental system where we could recapitulate normal infection progression with the ability to directly study host immunity.

We have developed a precision cut lung slice (PCLS) *ex vivo* model of cryptococcal infection. PCLS are an *ex vivo* tool with exceptional strengths for understanding host pathogen interactions that have yet to be properly exploited in the context of infection (17). Using PCLS in a *C. neoformans* murine infection model, we demonstrate that the early immune response involves limited phagocytosis and increased numbers of alveolar macrophages. We also examined other cell types, but found limited infiltration into alveoli by circulating myeloid cells. Alveolar macrophages are poor phagocytes of *C. neoformans* and phagocytosed cryptococci are able to grow efficiently within macrophages. In the lung, we found increased pathogen cell sizes, consistent with the early stages of titan cell formation. Finally, using the *C. neoformans plb1* mutant, we demonstrated the early differences that contribute to the avirulence of this mutant.

## Results

### Cryptococcal replication and clearance are balanced during the first 24 hours of infection

We found there was no change in the multi-lobe right lung homogenate fungal burdens in the first 24 hours following infection in a wild-type C57BL/6 mouse intranasal infection model using the highly virulent strain of *Cryptococcus neoformans*, KN99α GFP (18) (Fig 1A). We confirmed that the inoculum animals received at all time points were identical (Fig S1A) and observed no clinical signs of infection in infected animals (Fig S1B). Using our *ex vivo* infection precision cut lung slice (PCLS) model, we examined the distribution of dosed particles within the lung (Fig 1B). At a dose of 5×10^4^ cfu, there were typically no more than one cryptococcal cell per infected alveolus. We also used confocal microscopy to assess the appearance PCLS and found an exceptional preservation of lung anatomy, including the presence of alveolar macrophages (Fig 1C). As slices were taken from different animals and at slightly varying depths in the tissue, we also confirmed that PCLS were consistent in appearance (Fig S2). To confirm the sensitivity of the PCLS technique, mice were inoculated with different inert-polystyrene bead doses to confirm that the number of particles observed in PCLS correlated with differences in inoculum (R^2^=0.82) (Fig 1D). Therefore, we predicted we would see the same number of cryptococci in PCLS taken at 0 and 24hpi as fungal burdens were identical. Interestingly, however, we found significantly higher numbers of cryptococci in slices taken at 24hpi vs those taken immediately following inoculation (Fig 1E). This suggests that in specific regions of the lung, cryptococcal cells survive and replicate while in other areas there must be fungal clearance to explain the identical fungal burdens. There were no differences in burden or particle numbers when non-opsonised cryptococci and cryptococci opsonised with the highly specific 18B7 antibody were dosed (Fig 1A, D).

**Fig 1.**
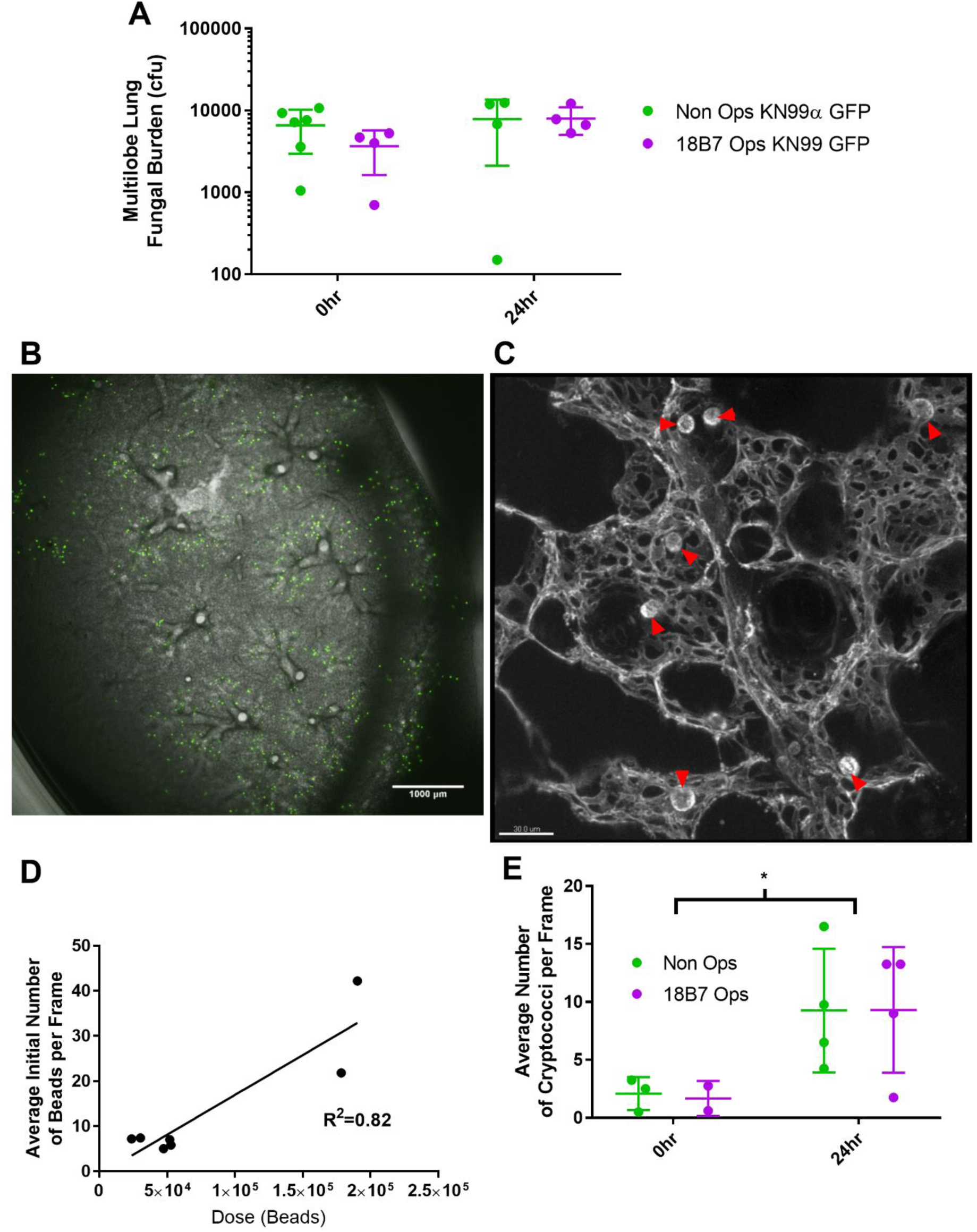
Precision cut lung slices reveal unchanged fungal burdens but increased fungal particles in specific lung loci. (A) Multi-lobe right lung fungal burdens at 0hpi and 24hpi (2 way ANOVA, Time: p=0.1485, Opsonin: p=0.4612, Interaction: p=0.4111)(n≥4). (B) Wide field image of precision cut lung slice and fluorescent beads (GFP) (C) confocal image of uninfected isolectin GS-IB_4_ stained PCLS, captured in Glasgow. Alveolar macrophages are indicated by red arrowheads. (C) Correlation between beads counted in PCLS and dose given to animals. Linear regression (n=7) (D) Number of cryptococci counted in PCLS at 0 and 24hpi. Two way ANOVA (Time: p=0.0186)(n≥2). Each point represents a different animal processed on different days.

### Macrophages are the primary cell population present during cryptococcal infection 24 hours post infection

In PCLS taken immediately following infection (0hpi), we labelled tissue with fluorescent isolectin GS-IB_4_, a plant dye that binds to both endothelial cells and alveolar macrophages (confirmed with CD11c counter staining; Fig 2A). Alveolar macrophages were dispersed throughout PCLS (Fig S3) at a density of less than one macrophage per alveolus (Fig 2B). There were consistent total numbers of alveolar macrophages across treatment groups, with numbers unaffected by the presence of 6μm beads or antibody opsonin (Fig 2C). At 24hpi, we used an antibody staining panel to attempt to identify interstitial dendritic cells, neutrophils, infiltrating monocytes and resident alveolar macrophages (Fig 2D-G) (19). All four cell types were present at 24hpi, although alveolar macrophages were the most abundant cell type in all cases. Again, there were no differences in cell numbers or types with bead and fungal inoculated animals, and antibody opsonin had no effect.

**Fig 2.**
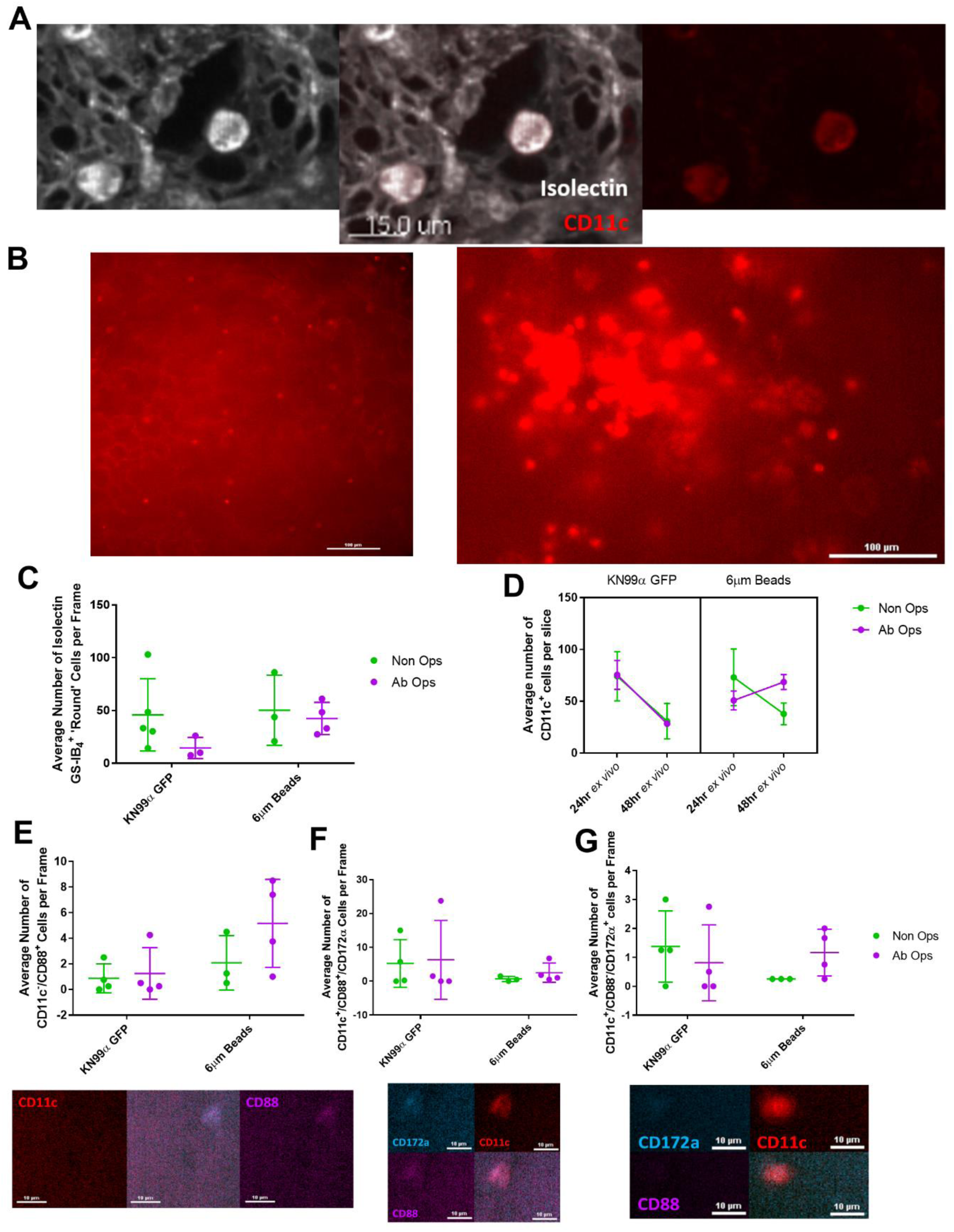
Immune responses at 0 and 24hpi to *Cryptococcus neoformans* KN99α and 6μm beads are alveolar macrophage centric. (A) Confocal image of uninfected PCLS showing colocalised CD11c FITC and Isolectin GS-IB_4_ AF-647, captured in Glasgow. (A) Left – Representative image of 0hpi PCLS with macrophages and blood vessels stained with Isolectin GS-IB_4_. Right – Representative image of 24hpi PCLS with CD11c AF647 stain (B(C) Number of isolectin GS-IB_4_ cells in 0hpi PCLS. (2 way ANOVA, ns). (D) Number of CD11c^=^ cells in 24hpi PCLS.(2 way ANOVA, Time=0.0003, interaction=0.003) (E) Top - Number of CD11c^−^/CD88^+^ cells in 24hpi PCLS. (2 way ANOVA, Time=0.0597). Bottom – Representative image of neutrophil (F) Top - Number of CD11c^+^/CD88^+^/CD172a^+^ cells in 24hpi PCLS. (2 way ANOVA, ns). Bottom – Representative image of monocyte derived macrophage (G) Top - Number of CD11c^+^/CD88^−^ /CD172a^+^ cells in 24hpi PCLS (2 way ANOVA, ns). Bottom – Representative image of dendritic cell. Each point represents a different animal processed on different days.

### Alveolar macrophages demonstrate limited phagocytosis of cryptococci

Having quantified the size of the immune presence in the lung, we next examined the exact nature of the host-pathogen interactions in PCLS at each time point, with a focus on alveolar macrophages. In PCLS prepared at 0hpi, macrophages showed minimal migratory behaviour and no incidences of phagocytosis were observed over 24 hours *ex vivo* across all four treatment groups. To confirm this wasn’t a technical limitation with our model, animals were instilled *ex vivo* with 10^9^ 1μm and/or 10^8^ 10μm beads, to confirm that phagocytosis in PCLS was preserved (20). We observed that phagocytosis of 1μm beads occurred rapidly before PCLS were mounted for imaging, and further phagocytosis was observed *ex vivo* (Fig 3A). However, despite the high inoculum of 10μm beads, we only observed one incidence of phagocytosis across 6 animals. The high rates of phagocytosis with 1μm beads *ex vivo* but very low to absent levels of phagocytosis with 6 and 10μm beads highlight the size dependent nature of phagocytosis. This suggests that the size of *C. neoformans* is inhibitory for phagocytosis (Fig 3B). Intriguingly, 24hpi PCLS contained phagocytosed cryptococci, indicated that there are *in vivo* differences in phagocytosis, independent of antibody opsonisation (Fig 3C-F).

**Fig 3.**
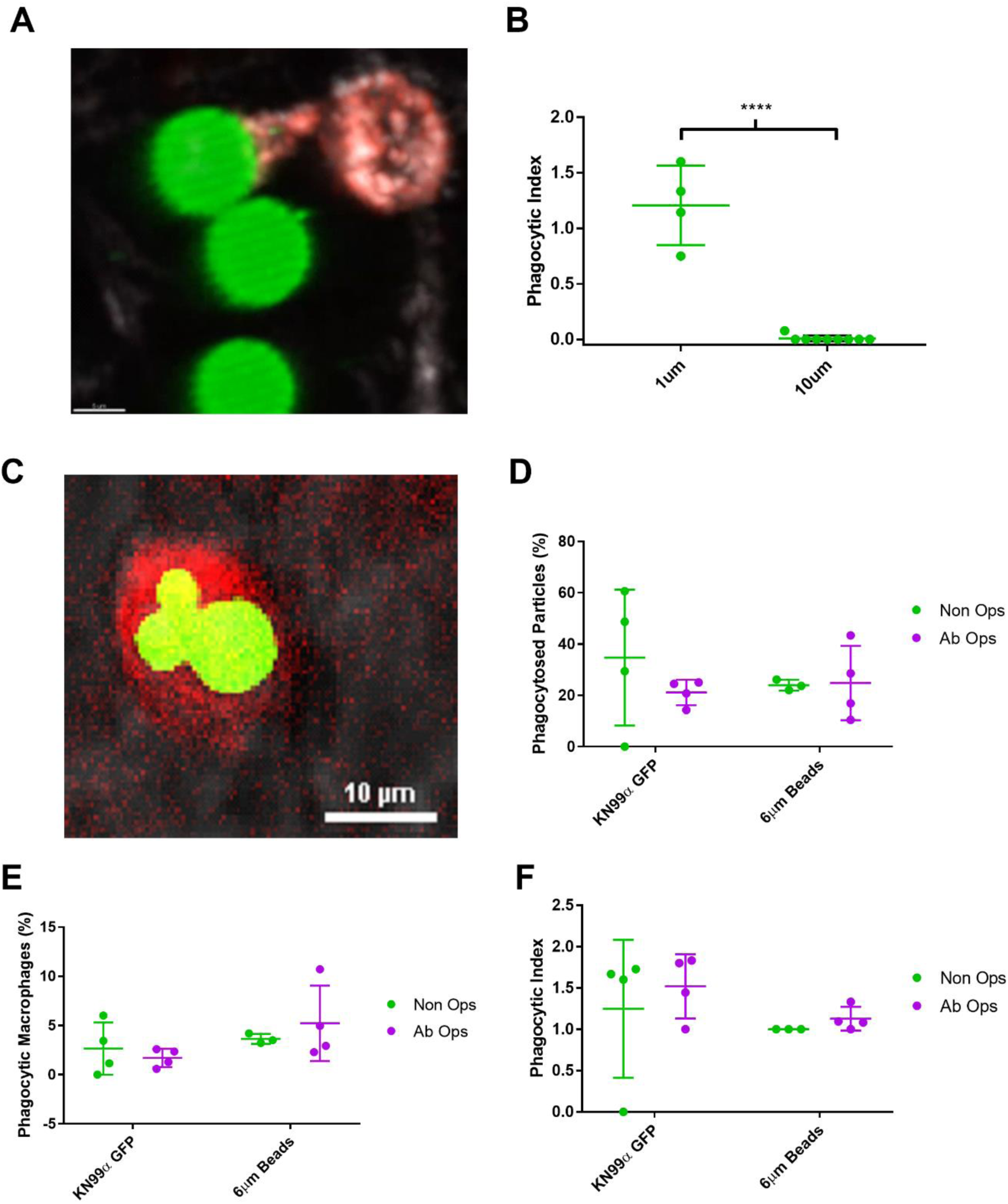
Phagocytosis of *Cryptococcus neoformans* KN99α GFP and 6μm beads only occurs at 24hpi, not 0hpi. (A) Representative image of a 1μm bead, not a 10μm bead, being phagocytosed by an alveolar macrophage (Isolectin GS-IB_4_ – white, CD11c – red) (n=1 animal, 2 videos) (B) Phagocytic index of 1μm and 10μm beads in PCLS (unpaired two tailed t-test, p<0.0001) (C) Representative image of phagocytosed *Cryptococcus neoformans* in 24hpi PCLS (D) Percentage of particles phagocytosed within 24hpi PCLS (2 way ANOVA, ns). (E) Number of macrophages that phagocytosed particles in 24hpi PCLS (2 way ANOVA, ns). (F) Phagocytic index of particles in 24hpi PCLS (2 way ANOVA, ns).

### Cryptococcal replication inside macrophages was indistinguishable from growth extracellularly

Having assess immunity to *C. neoformans*, we then examined the phenotype of cryptococcal cells. We observed cryptococcal budding both in 0hpi and 24hpi PCLS (Fig 4A-B). Interestingly, the number of cryptococcal particles in 0hpi PCLSs after 24 hours of culture *ex vivo* was the same as the number of cryptococci in 24hpi PCLS (Fig 4C). This suggests a strong correlation between conditions in the lung *in vivo* and our *ex vivo* model.

**Fig 4.**
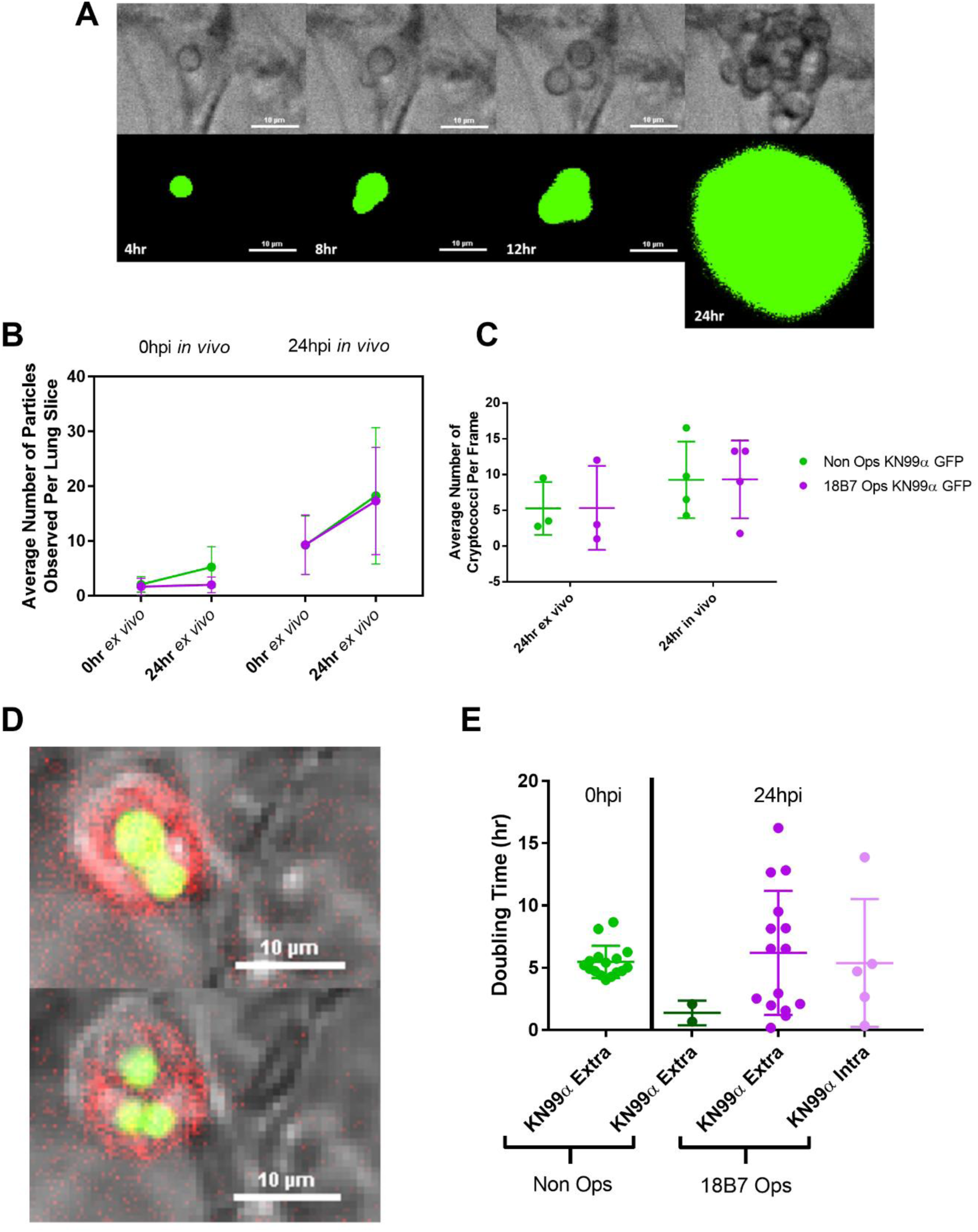
*Cryptococcus neoformans* KN99α GFP replicates *ex vivo* in PCLS. (A) Representative images of cryptococcal replication within 0hr PCLS. (B) Change in number of cryptococcal cells in 0hpi and 24hpi PCLS (2 way ANOVA, Time=ns). (C) Comparison of number of cryptococci in 0hpi PCLS after 24hr imaging *ex vivo* and 24hpi PCLS (2 way ANOVA, ns). (D) Representative image of cryptococcal intracellular replication. (E) Comparison of average doubling times of cryptococci in different conditions in both 0hpi and 24hpi PCLS. Each point represents the growth characteristics of a single cryptococcal cell (1 way ANOVA, ns).

One of the major features *of C. neoformans* is thought to be the ability to grow intracellularly inside macrophages. This has been previously hypothesised based on histology in a mouse model of cryptococcosis and *in vitro* observations (21, 22). Therefore, we wanted to confirm if this also occurred in our *ex vivo* model where individual cells could be tracked. We observed that phagocytosed cryptococcal cells replicated *ex vivo* (Fig 4D). Then, to understand this growth further, we derived average growth equations for each condition, to see if the macrophage niche was permissive or inhibitory for growth. We found that the growth of *Cryptococcus* across all conditions did not significantly differ, including intracellularly (Fig 4E), suggesting that phagocytosing *Cryptococcus* did not limit fungal growth.

### Cryptococcal cell size increases early in infection

Another of *C. neoformans* pathogenic strategies is the ability to form cells with massively increased size *in vivo* (>30μm), described as ‘titan cells’ (13). As this phenotype has only just been recapitulated *in vitro*, our understanding of this phenotype *in vivo* is still poorly understood (23). In our model, a variety of cryptococcal cell sizes were observed, but no consistent differences in size were observed between 0hpi and 24hpi. The requirement for a cell to be classed as a titan cell is a total diameter of >30μm and although no cells this extreme were found, we did identify several cells with a total diameter >10μm (Fig 5C-D) (24).

**Fig 5.**
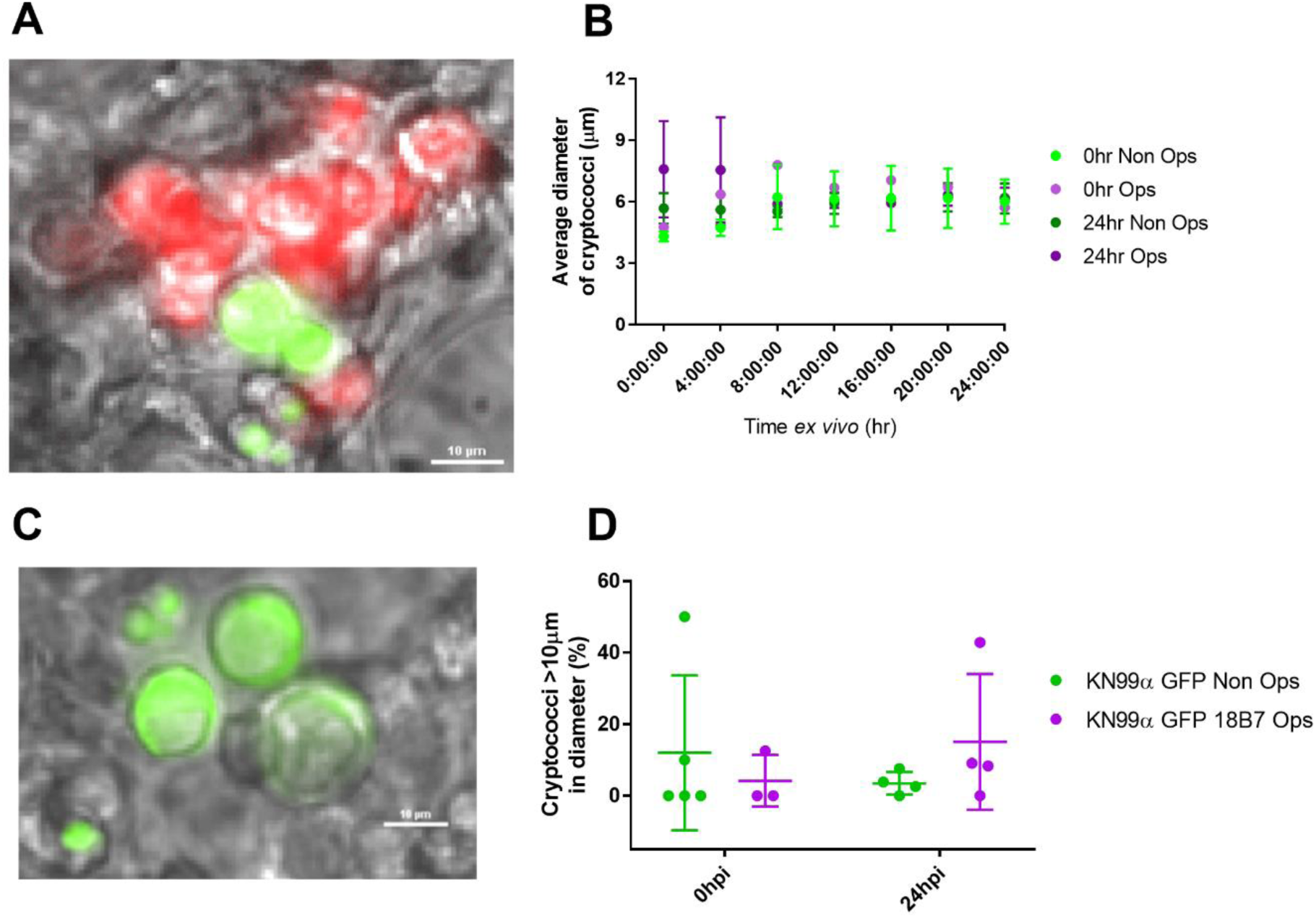
*Cryptococcus neoformans* KN99α GFP does significantly change size between 0 and 24hpi or *ex vivo*. (A) Representative image of different cryptococci sizes in 24hpi PCLS. (B) The average size of cryptococcal cells over 24hr *ex vivo* in 0 and 24hpi PCLS (2 way ANOVA with repeated measures, ns). (C) Representative image of large KN99α GFP cryptococcal cells observed in 24hpi PCLS. (D) Proportion of KN99α GFP cells larger than 10μm in diameter (2 way ANOVA, ns).

### plb1 *mutant* C. neoformans *exhibit increased cell size and are not phagocytosed*

Having characterised the highly virulent *KN99α* strain in the PCLS model, we examined an avirulent strain of *C. neoformans* to identify any differences in pathology within the first 24 hours of infection or any clear differences in immunity. We selected the *plb1* H99 GFP strain of *C. neoformans*, known to cause no mortality in the mouse model and known to be unable to replicate inside macrophages (25, 26). We found no significant difference in mouse clinical signs (Fig S3), multi-lobe right lung homogenate fungal burden (Fig 6A) or in the number of fungal cells 24hpi (Fig 6B) with *plb1* inoculated animals compared to *KN99α* inoculated animals. Importantly, there were no differences in leukocyte numbers with either *KN99α* or *plb1* infection (Fig 6B-E). Despite an identical number of leukocytes, we observed no phagocytosis of *plb1* mutant cells at 24hpi (Fig 6F-G). Given these findings, we then investigated whether *plb1* mutant cells demonstrated any phenotypic differences that may explain the heightened resistance to phagocytosis. There was no difference in extracellular fungal replication (Fig 7A-B), we did observe that *plb1* cells were significantly larger *ex vivo* compared to the highly virulent KN99α GFP cells. This similar to what has been observed both *in vitro* and later in infection *in vivo* (Fig 6C) (26). The proportion of *plb1* H99 GFP cells larger than >10μm in diameter was significantly higher, suggesting that the *plb1* mutant may be more predisposed to form titan cells (Fig 6D).

**Fig 6.**
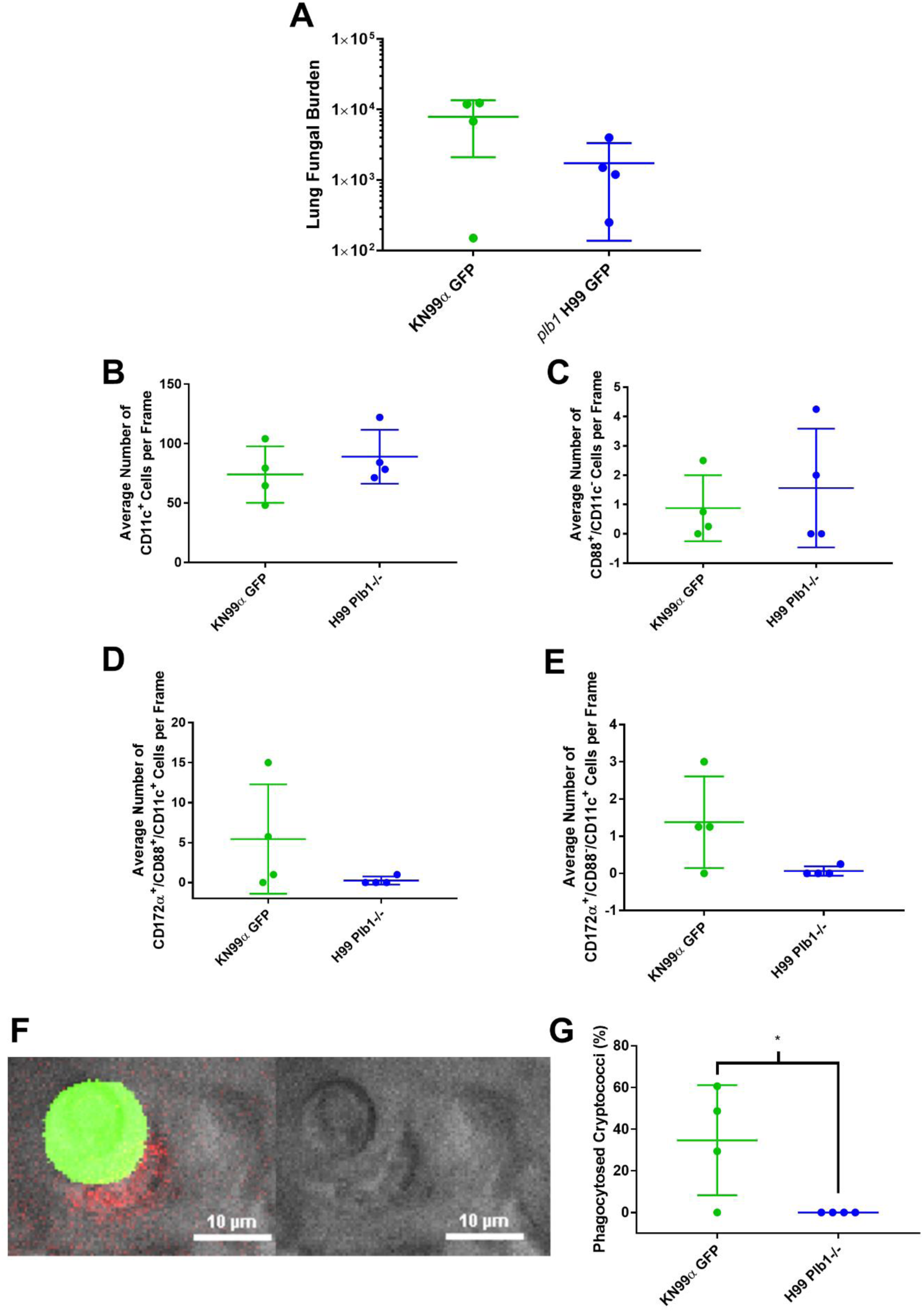
*Cryptococcus neoformans plb1* H99 GFP cells are significantly larger than KN99α GFP when cultured over 24hr in 24hpi PCLS. (A) Multi-lobe right lung fungal burden at 24hpi for KN99α GFP and *plb1* H99 GFP (unpaired 2 tailed t-test, p=0.086). (B) Number of particles in 24hpi PCLS (unpaired 2 tailed t-test, ns). (C) Proportion of each strain of *Cryptococcus neoformans* that replicated in 24hpi PCLS (unpaired 2 tailed t-test, ns). (D) Average size of KN99α GFP and *plb1* H99 GFP in 24hpi slices (2 way ANOVA with repeated measures, Time=0.0017, Strain=0.0012, Interaction=0.0307. *post hoc* Sidak multiple comparisons left to right 0.0038, 0.0013, 0.0005, 0.0008, <0.0001, <0.0001. (E) Proportion of KN99α GFP and *plb1* H99 GFP cells in 24hpi slices that are larger than 10μm in diameter (unpaired two tailed t-test, p=0.0417).

**Fig 7.**
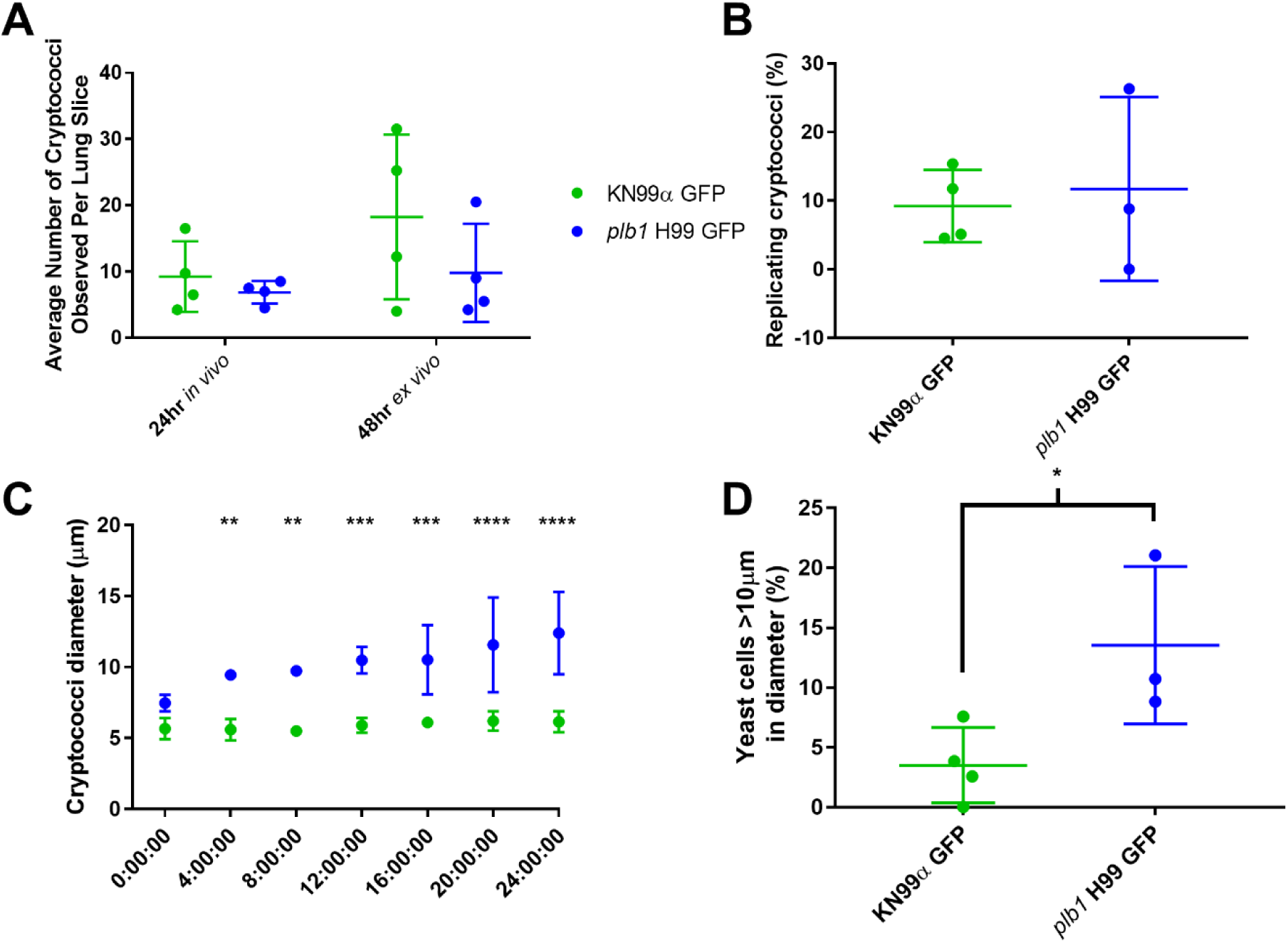
The immune response to KN99α GFP and Plb1^−/−^ H99 GFP does not differ in 24hpi PCLS. (A) Number of CD11c^+^ positive cells in 24hpi PCLS (unpaired 2 tailed t-test, ns). (B) Number of CD88^+^/CD11c^−/−^ cells in 24hpi PCLS (unpaired 2 tailed t-test, ns). (C) Number of CD11c^+^/CD88^+^/CD172a^+^ cells in 24hpi PCLS (unpaired 2 tailed t-test, ns). (D) Number of CD11c^+^/CD88^−^/CD172a^+^ cells in 24hpi PCLS (unpaired 2 tailed t-test, ns). (E) Representative image of failed phagocytosis of *plb1* H99 GFP by an alveolar macrophage. (F) Proportion of cryptococci phagocytosed in 24hpi PCLS (unpaired 2 tailed t-test, p=0.0395).

## Discussion

Here, by combining *in vivo* mouse infection with an *ex vivo* PCLS model of cryptococcosis, we present the first visualisation of the pulmonary immune response to *C. neoformans* in the first 24 hours of infection and show it is not sufficient for clearance. At the cellular level, resident alveolar macrophages appear to be the predominant cell type present at sites of infection. Phagocytosis does not occur immediately following inhalation of fungal cells and only occurs at a low frequency during the first 24 hours of infection. However, even if phagocytosis does occur, alveolar macrophages appear to provide a permissive environment for intracellular growth. This suggests that early cell mediated immunity isn’t sufficient to clear cryptococcal infection. Cryptococci became enlarged early in infection, a phenotype that is more pronounced in the *plb1* mutant. The attenuated *plb1* mutant is able to grow extracellularly in the lung, but critically appears resistant to phagocytosis.

Despite the fact *C. neoformans* is an opportunistic infection, we failed to see a reduction in burden over 24 hours of infection, as has been seen previously in studies examining *KN99α* fungal burdens 1dpi (27). This is likely an effect of inoculating animals with 50,000cfu, a higher dose than environmental exposure would likely ever be. This widely published dose corresponds to no more than one cryptococcal cell per alveoli, while being tractable for live imaging (28). However, at this dose the immune response is likely unable to respond to all yeast cells rapidly enough for clearance. This delay allows surviving fungal cells to replicate, grow in size and establish host evasion mechanisms, highlighting the stochastic nature of opportunistic infection. Our observation that there is a balance of cryptococcal clearance and replication during this first 24-hour period means there are several factors that dictate which event predominates. By understanding these processes and identifying critical signalling pathways, we may find new factors that enhance cryptococcal clearance or inhibit replication.

The macrophage-centric response to *Cryptococcus neoformans* has long been hypothesised, due to the number of macrophages present in multiple clinical observations of cryptococcosis (29). However, this is the first direct evidence that alveolar macrophages are most abundant cells at sites of *C. neoformans* infection this early in infection (30). The minimal recruitment of infiltrating cell types, however, suggests that *C. neoformans* is not inflammatory enough to cause the mass infiltration of leukocytes 24hpi (31). Alternatively, recruitment may have occurred between 0 and 24hpi, and subsequently reverse migration/apoptosis. Therefore, characterising the immune response at 4-6hpi is still required to accurately plot the timeline of the response. The low number of neutrophils present is also unsurprising. Neutrophils are a particularly short-lived cell type (<24 hours) (32) and it is possible that these cells would no longer be present at the site of the infection if recruited early in infection. Furthermore, there is currently no clinical association between neutrophil disorders and incidences of cryptococcal disease, which implies that these cells are not critical during cryptococcal infection. Dendritic cells, on the other hand, are known to be resident in the lung and have been shown to have antifungal activity (33). However, we saw limited numbers during our study and those we did observe did not directly interact with cryptococci, in contradiction to previous work. It is important to note that differentiating between infiltrating and resident macrophages is notoriously difficult (34). Although several studies have suggested surface markers that differentiate the two, there is still a level of uncertainty with our immunofluorescence approach and should be supported by alternative methodologies (19, 35). The other question raised about immunity to *Cryptococcus* in our model is why we did not directly observe or recognise fungal killing *ex vivo*, despite evidence that clearance occurred in the first 24 hours of infection. One explanation is that nutrient starvation is hypothesised to be one mechanism by which macrophages kill phagocytosed pathogens. In our PCLS methodology, we may have inadvertently affected this process as we use a relatively glucose rich media to maintain viable slices (6).

There was a complete lack of a host or pathogen phenotype in the presence of antibody opsonin. This contradicts many *in vitro* cryptococcal studies, where 18B7 antibody is said to be required for phagocytosis by macrophage and macrophage like cell lines to occur (36). Our work presented here highlights that exclusively modelling phenomena *in vitro* using cell lines does not accurately recapitulate the *in vivo* environment, meaning that results may be artificial and not representative of *in vivo* infection. Mechanistically, the lack of enhanced phagocytosis at 24hpi suggests a minimal role of the Fc receptor until later in infection. This should be confirmed using a different opsonin, such as complement protein, which may be a more critical factor for early phagocytosis.

Titan cells, a common clinical observation (37), are difficult to generate *in vitro*. We did not observe titan cell formation in our model, but as the recently established *in vitro* protocol for titan cell generation takes 3 days, this suggests this phenotype does not occur this early in infection (23). This is supported by *in vivo* observations, where mice are only examined *in vivo* for titan cell formation at 3dpi (24). Therefore, cells may take longer than this to reach this phenotype. Despite no titan cells, we did find cryptococcal cells more than doubled in diameter during our experiments, potentially approaching a titan cell phenotype.

The *plb1* mutant is well characterised, known to produce larger cells, is unable to disseminate and is unable to replicate within macrophages (26). The major difference in the immune response to these two strains was that we observed no phagocytosed *plb1* mutant cells 24hpi. This implies that despite the mutant strains inability to produce immunomodulatory compounds (PGE_2_), the significantly larger cells overcompensate, making cells extremely resistant to phagocytosis later in infection (26). However, this result provides conflicting conclusions. The inability of these cells to be phagocytosed suggests the *plb1* mutant cryptococcal cells are better at immune evasion and so less likely to be cleared by the immune system and better at colonising the host. However, previous work has shown that *plb1* lung burdens at 7dpi are significantly lower than time-matched virulent strain burdens (38). The same study also noted lower total lung leukocyte numbers, indicative of a dampened immune response. This suggests that being resistant to phagocytosis and evading immunity is actually detrimental to the pathogen and that the macrophage intracellular niche in important for replication and cell survival. Dissemination also appears to be dependent on macrophages, as *plb1* infected mice showed no brain burden, even at 3 weeks post infection. Combined, there is strong evidence that alveolar macrophages are the ‘villain’ in cryptococcosis, providing a protective niche while aiding cryptococcal dissemination via a ‘Trojan horse’ mechanism (39).

The model used in this study - PCLS - are extremely resilient and easy to maintain, reported to remain viable for up to 7 days, allowing for long term investigation (17). This *ex vivo* approach also increases the number and type of experimental measures available during *in vivo* studies while simultaneously reducing the need to subject animals to procedures *in vivo*. Furthermore, as PCLS is a relatively simple methodology associated with few initial costs, it is possible for most laboratories to adopt this technique; an important factor in its utility in reducing animal use. PCLS is also a highly flexible technique that can be easily modified for purpose. Here we have focussed on high-resolution time-lapse imaging, but also demonstrate how it is possible to image slices at a lower magnification and assess larger scale phenomena, as well as lung structures other than the alveoli. A single mouse lung can yield 10+ individual slices, enabling correlative techniques, e.g. –omics approaches, high-content screening approaches such as targeted therapeutic identification. This model is also not specific to cryptococcal infection so can be used with any pulmonary pathogen, providing a great opportunity to study polymicrobial infections. Finally, this technique is also not limited to the lung, and has also been adapted for use with other organs, such as the liver (40).

However, it should be noted that PCLS does have some limitations. Although most physiological processes, such as immunity and airway contraction, are preserved *ex vivo* (41), circulation is completely absent. This may greatly affect studies of various phenomena, in particular dissemination of *C. neoformans* out of the lung or directly visualising the extravasation of circulating immune cells. However, other models may be appropriate in this instance (e.g. long-term intravital imaging in the zebrafish). Alternatively, conditions in PCLS may be correlated with short-term intravital imaging in mouse and/or brain fungal burdens. Another downside is that once a slice has been prepared for PCLS, there is no outside recruitment of cells. While this may be beneficial for the study of resident cells in isolation, this does limit the contexts in which this method is appropriate. Finally, it should be noted that although fungal burden analysis only detects viable cells, imaging of PCLS potentially included non-viable and/or killed fungal cells, which may have skewed results. Therefore, a mutant cell line that indicates cell viability should be utilised to allow viable and dead cells to be distinguished visually.

To conclude, for the first time we directly visualise the innate immune response to the opportunistic pathogen *Cryptococcus neoformans*. We examined the different cells present and identified alveolar macrophages as the major cell type present as sites of infection and that these cells are insufficient to clear infection. We also observed fungal size increases, as well as the first recorded instance of cryptococcal replication within the lung. We believe this approach is highly tractable and will be widely adopted to study infection and immunity in the future.

## Methods

### Ethics Statement

Animal work was carried out in accordance with both national and institutional ethical guidelines. All procedures were carried out by competent individuals in accordance with the Home Office Animal (Scientific Procedures) Act 1986, under project licenses 40/3726 or P4802B8AC after ethical review by the Animal Care and Welfare Committee of the University of Sheffield.

### Mouse Infections

C57BL/6 mice aged 8-13 weeks were purchased from Charles River (UK) and were housed in The University of Sheffield animal unit, except for *ex vivo* 1 and 10μm bead comparison experiments, which were performed on humanely killed 8-13 week old C57BL/6 mice that were housed at the Beatson Biological Research Unit. Mice were randomised into treatment groups using the Google random number generator. Animals were infected by intranasal inoculation - mice were lightly anaesthetised using gaseous isofluorane and inoculated with 50μl of either 50,000cfu *Cryptococcus neoformans* KN99α GFP (18) or H99 ΔPlb1 GFP (42, 43) or 50,000 6μm Fluoresbrite® YG Microspheres Polysciences (Hamburg, Germany). 0hpi mice were sacrificed by pentobarbital injection and exsanguination following recovery from the intranasal dosing anaesthesia. 24hpi mice were individually housed in individually vented cages once inoculated, and weighed daily to monitor health status.

### Cryptococcus culture, preparation and opsonisation

Unless stated, all reagents were purchased from Sigma (Dorset, UK). Prior to use, *Cryptococcus* strains were cultured for 18 hours in YPD medium. Cultures were rotated continuously at 30rpm and kept at 28°C on a 14hr:10hr light/dark cycle. Cultures were washed by centrifuging cultures (6,000rpmrpm, 1 minute) in PBS before being counted and diluted to the desired concentration in PBS. For opsonised *Cryptococcus neoformans* KN99α GFP, mouse 18B7 antibody, an IgG antibody against cryptococcal capsule, was applied to counted cultures at a dilution of 1:1000 for one hour at 37°C (44, 45).

### Bead preparation and opsonisation

Beads were prepared identically to cryptococcal cultures, with PBS washes followed by counting. In order to opsonise the beads, 10^8^ were incubated overnight at 4°C in PBS containing 10mg/ml BSA. Samples were washed in PBS before being opsonised with mouse anti-BSA antibody at a dilution of 1:1000 for an hour at 37°C. Opsonised beads were prepared ahead of time and were stored until use away from direct sunlight at 4°C.

### Lung slice preparation

Immediately after humane killing by i.p. pentobarbital overdose and exsanguination, the mouse trachea was exposed by blunt dissection and cannulated. Warm 2% low melting point agarose was instilled into the trachea to inflate the lungs. Cadavers were placed on ice for ∼2 minutes to allow the agarose to set. Once set, the lungs were carefully removed from the animal and suspended in PBS. The multi-lobe right lung was taken for burden analysis and the single lobed left lung was attached with tissue adhesive (Vetbond, IMS (Euro), Stockport, UK) to a cutting block, and was then placed in a Vibratome (Leica, London, UK). The lung was bathed in high glucose DMEM supplemented with 10% FBS and 1% penicillin-streptomycin. 300μm thick slices were then prepared and high quality slices were mounted in an imaging dish, and weighed down using stainless steel wire and thin fishing wire to counteract the buoyancy of the tissue. For Glasgow bead comparison experiments, PCLS were weighed down using Tissue Slice Anchors (Warner Instruments, Connecticut, USA).

### Antibody Labelling

For 0hr experiments isolectin GS-IB_4_ conjugated to AF647 (ThermoFisher, Loughborough, UK) was used to visualise blood vessels and alveolar macrophages. The dye was added into the imaging well at a dilution of 1:200. For 24hr experiments, the multicellular response anticipated meant that a variety of antibodies were used to capture the major innate cell populations. Antibodies were selected according to Lyons-Cohen et al. (19). CD88 (BioRad, Kidlington, UK) and CD172α (Biolegend, London, UK) antibodies were conjugated to AF 594 and AF555 respectively using conjugation kits according to supplier instructions. CD11c antibody conjugated to AF647 (Biolegend, London, UK) was also used. CD11c was added to the imaging well at a dilution of 1:200, whereas CD172α and CD88 conjugates were added at a dilution of 1:50.

### Imaging

Long term 24hr imaging was done using a widefield Nikon Ti (Kingston Upon Thames, UK), imaged at 20x using the settings, lens and camera described in Evans et al. (43). All data was captured using NIS element AR software. The Z axis was controlled using a piezo motor. For confirmation of the isolectin/CD11c staining and bead comparison experiments, an LSMZeiss 880 Airyscan Fast confocal micscope with 20× 0.8 N.A. air objective (Cambridge, UK) microscope was used in Airyscan Fast mode with default processing options in Zeiss Zen Black software.

### Image Processing and Analysis

All widefield images were visualised and analysed in .nd2 format in NIS Element Viewer (Kingston Upon Thames, UK) or ImageJ. Confocal images were visualised using Imaris software (Oxford Instruments). For cryptococcal size changes, individual cryptococci were manually segmented and measured using NIS Elements AR software. For cryptococcal growth equations, budding was tracked and used to derive growth equations for individual cryptococci using a non-linear fit of exponential growth in GraphPad Prism 7. Equation parameters for each treatment group were averaged to derive an average growth equation for each group. For size changes, 5-10 cryptococcal cells were randomly sampled and their diameter was measured manually using NIS Elements. Cells were measured in the GFP channel.

### CFU Fungal Burden Analysis

The multi-lobe right lung was weighed and then suspended in 500μl PBS. The tissue was then homogenised using a tissue homogeniser (Cole Parmer, Illnois US). The homogenate was then serially diluted from 1:2 to 1:20000 in PBS. 10% YPD agar plates were prepared, and 20μl homogenate was plated out. Plates were cultured for 48 hours at 28°C, where cultures were counted.

### Statistical Analysis

For all comparisons at the same time point between the 4 groups, 2 way ANOVA with *post hoc* Tukey comparisons were carried out. Growth curves were generated using the GraphPad exponential growth equation feature. Only equations with 3 points or more, and a R^2^ >0.7 were considered for analysis. Each point presented represents a single animal. Each animal was used on a separate day, and so counts as a biological repeat. All analysis was done using GraphPad Prism 7 (San Diego, USA) Statistical significance was taken to be p<0.05.

## Acknowledgements

We thank the Renshaw and Elks labs for critical discussions. We also thank the Sheffield and Glasgow animal unit staff, in particular Carl Wright, for their help with mouse husbandry and *in vivo* training. We are also grateful to Core Services and Advanced Technologies at the Cancer Research UK Beatson Institute (C596/A17196), with particular thanks to the Beatson Advanced Imaging Resource. JR is supported by the Medical Research Council DiMeN DTP (https://www.dimen.org.uk/) and Medical Research Council Flexible Funding. SAJ was supported by Medical Research Council and Department for International Development Career Development Award Fellowship MR/J009156/1 (http://www.mrc.ac.uk/). SAJ was additionally supported by a Krebs Institute Fellowship (http://krebsinstitute.group.shef.ac.uk/), and Medical Research Council Centre grant (G0700091). LMC is grateful for funding from Cancer Research UK (core funding A23983 and A17196). HMM is supported by funding from the Medical Research Council SHIELD consortium MRNO2995X/1.

