## Supplemental Figures for "The mouse lung early cellular innate immune response is not sufficient to control fungal infection with Cryptococcus neoformans"

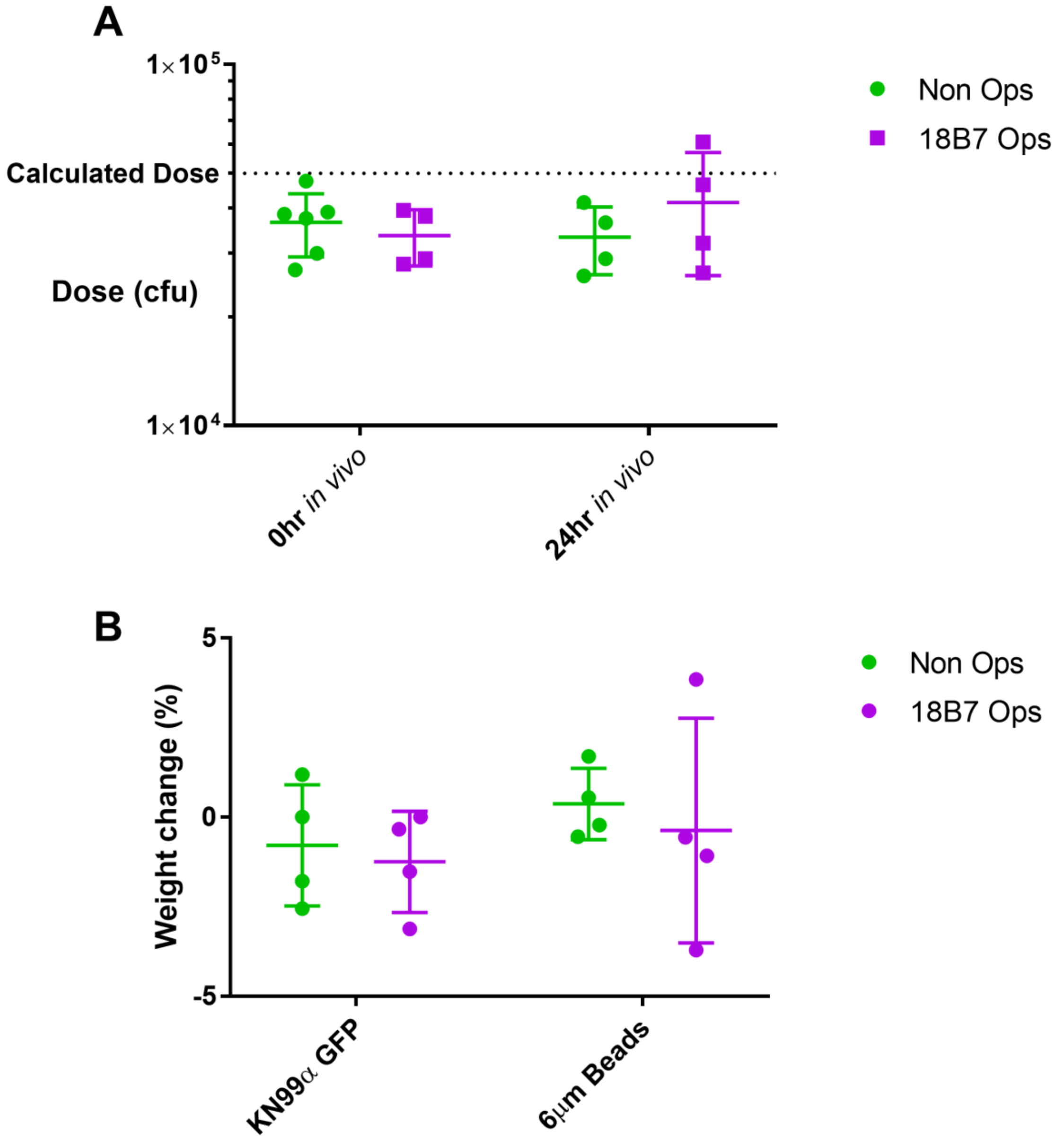


**Fig S1. Infection phenotype in KN99α GFP 24hpi animals**

(A) Actual doses of KN99 GFPα (2 way ANOVA, ns). (B) Weight change of C57BL/6 mice infected with KN99α GFP (2 way ANOVA, ns).

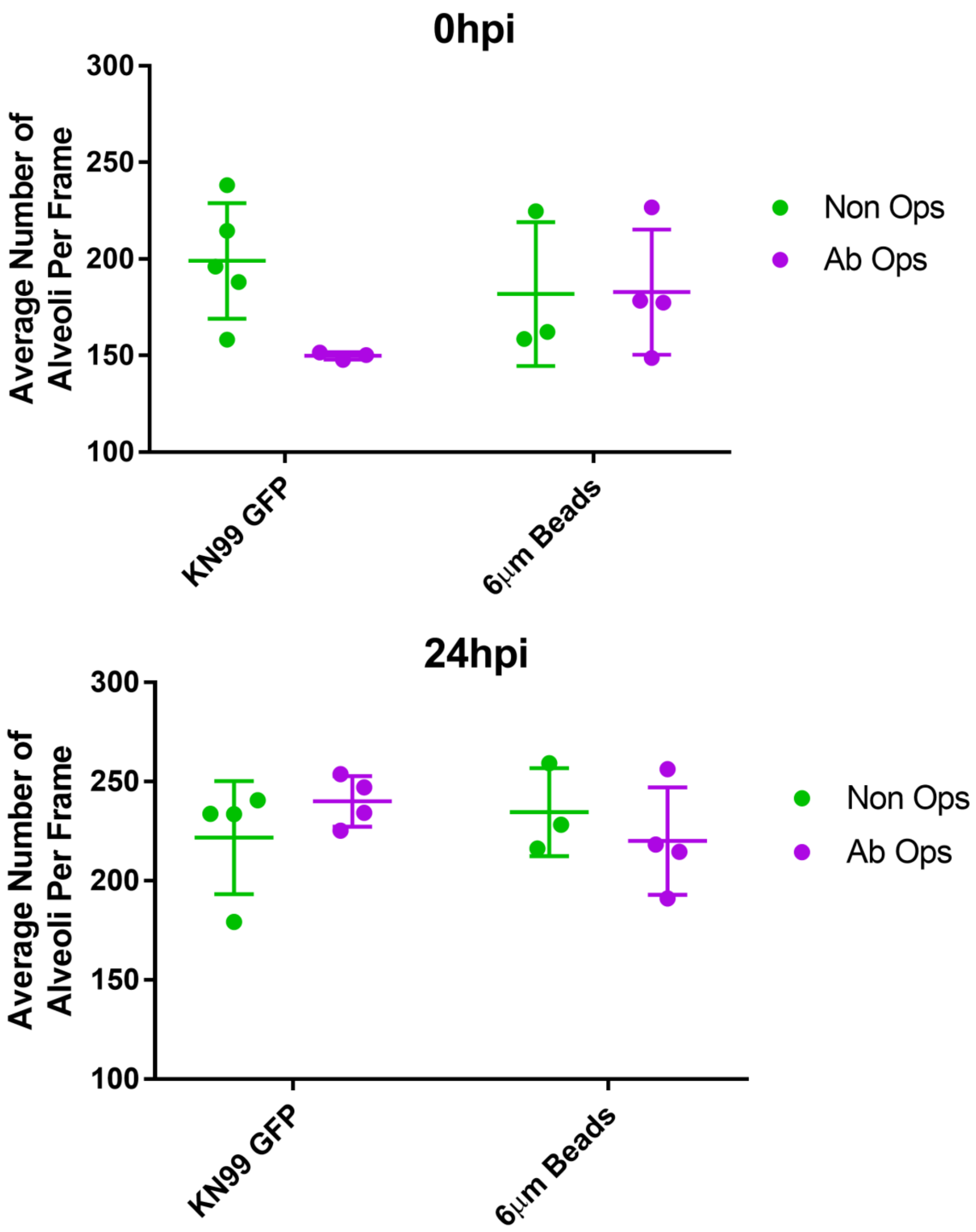


**Fig S2. Appearance of PCLS at 0 and 24hpi**

Number of alveoli counted in 0hpi PCLS (2 way ANOVA, ns) and 24hpi PCLS (2 way ANOVA, ns).

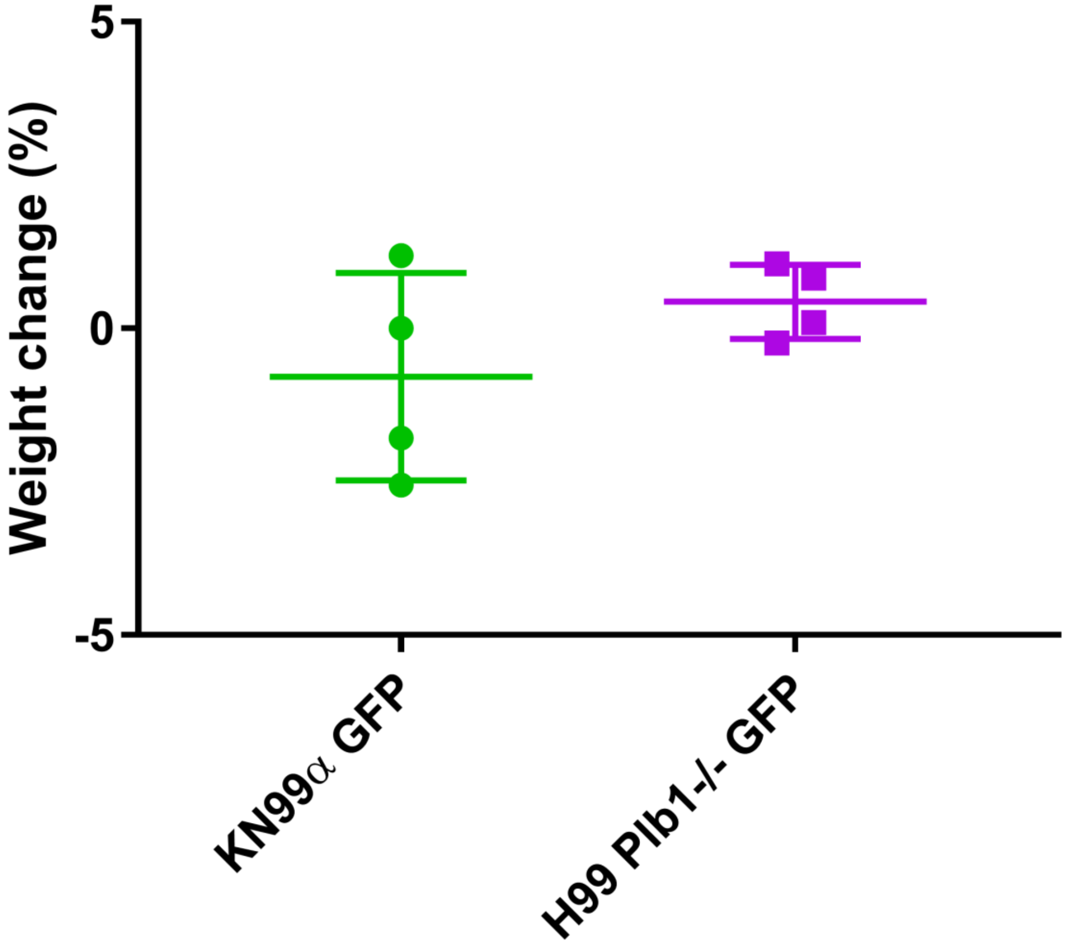


**Fig S3. Infection phenotype in KN99α GFP and *plb1 H99* GFP inoculated animals at 24hpi**

Weight change of animals inoculated with either KN99α GFP and *plb1 H99* GFP (2 way ANOVA, ns).
